# Acceleration of genome replication uncovered by single-cell nascent DNA sequencing

**DOI:** 10.1101/2022.12.13.520365

**Authors:** Jeroen van den Berg, Vincent van Batenburg, Alexander van Oudenaarden

## Abstract

In a human cell thousands of replication forks simultaneously coordinate the duplication of the entire genome. The rate at which this process occurs, might depend on the epigenetic state of the genome and vary between, or even within, cell types. To accurately measure DNA replication speeds, we developed a technology to detect recently replicated DNA using single-cell sequencing. Replication speed is not constant but increases during S-phase of the cell cycle. Using genetic and pharmacological perturbations we are able to alter this acceleration of replication and conclude that DNA damage inflicted by the process of transcription limits the speed of replication during early S-phase. In late S-phase, during which less transcription occurs, replication accelerates and approaches its maximum speed.

## Introduction

A large number of cell divisions are required to construct an adult animal body from a one-cell embryo. Before each cell division, the genome has to be faithfully replicated and DNA replication errors have to be averted to prevent developmental defects and tumorigenesis(1, 2). DNA replication can be roughly divided in four steps; origin licensing, origin firing, fork elongation and fork termination(3). Prior to DNA replication, the pre-replicative complex (pre-RC) is loaded on specific genomic regions, known as replication origins(4). The assembly steps of the pre-RC are tightly regulated and separated in time, which prevents multiple rounds of origin licensing in one cell cycle. After licensing DNA replication origins, the cell accumulates pro-proliferative signals until it reaches the threshold required to build the replication machinery (or replisome) at the pre-RC(4). Once the replication machinery is assembled at the pre-RC, several kinases are recruited and their stimulatory activity induces the firing of replication origin(5). However, not all licensed DNA replication origins fire simultaneously(6). Limiting origin firing is required to support the high demand of replisomes on both nucleotides and histones(7, 8). Following activation of the replisome, DNA helicases unwind the DNA double strand which allows subsequent copying of both strands by DNA polymerases(9). However, the copying process of both strands is not identical due to directionality of the DNA molecule; these different modes of replication are referred to as the leading and lagging strand synthesis(10). Finally, two converging replisomes result in the termination of DNA replication for that particular location in the genome. The replisomes get dissembled when in close proximity and residual replication is mediated by a complex hand-off between several DNA polymerases(11). The fields of genetics and biochemistry have revealed many DNA replication factors and how they cooperate to ensure high fidelity duplication of genomes. However, these methods are limited in their ability to probe the positional information of DNA replication. More recently, DNA sequencing methods have been used to unravel replication timing, replication fork directionality, origin and DNA polymerase usage(12–15). Especially, single-molecule approaches have been invaluable to monitor this process by visualizing DNA replication and subsequent detection by microscopy(16, 17) or long-read sequencing(18, 19). However, these methods randomly sample molecules from a large population of cells and are therefore insensitive to heterogeneity in replication dynamics between individual cells. Here, we describe a novel method, scEdUseq, which allows high resolution single-cell investigation of DNA replication fork dynamics.

## Results

### scEdU-seq reveals ordered DNA replication profiles throughout S-phase

In order to measure heterogeneity of DNA replication fork dynamics between cells, we developed single-cell EdU sequencing (scEdU-seq), a sequencing method to identify replicated nascent DNA in individual cells. scEdU-seq relies on metabolic labeling with the nucleotide analog 5-Ethynyl-2′-deoxyuridine (EdU) and subsequent affinity capture of newly synthesized DNA fragments (Figure 1A). We make use of CuAAC Click chemistry to covalently link a biotin moiety to the uracil base(20). Subsequently, we digest the single-cell genome and ligate adapters containing a T7 promoter, cell-specific barcodes and a unique molecular identifier (UMI)(21, 22). After pooling cells, we biotin-capture the EdU containing DNA molecules and release the non-EdU modified strand by heat denaturation. Finally, we regenerate the complementary strand via primer extension followed by amplification to prepare for sequencing. We first compare scEdU-seq to Repli-Seq(15) in human RPE-1 hTERT cells and find high overlap of early and late DNA replication profiles (Figure 1B, Supplemental Figure 1A,B). Next, we set out to generate DNA replication profiles in single cells (Supplemental Figure 1C,D). We can use these single-cell DNA replication profiles to reconstruct the progression through S-phase by ordering the cells based on the overlap coefficient(23) and by computing a 1-D UMAP(24), which we define as “S-phase progression” (Supplemental Figure 1E,F). We observe that the S-phase progression accurately reflects the position in the cell cycle as determined by the FUCCI(25) reporters, a fluorescent read-out which reflects cell cycle state, as well as DNA content (Figure 1C,D, Supplemental Figure 1G). Implementing the single-cell S-phase progression ordering, we construct DNA replication tracks over the entirety of S-phase from an ensemble of 1343 RPE-1 hTERT FUCCI cells (Figure 1E). The start of these tracks coincides with positions in the genome that were previously identified as initiation zones (IZ) of DNA replication in RPE-1 cells(26). Furthermore, our S-phase ordered scEdU-seq data is consistent with published bulk DNA replication timing using cell-cycle sorted populations(27). In addition, we observe that in late S-phase the scEdU-seq domains consist of large blocks of active DNA replication, whereas the middle of S-phase mainly consists of long-lived replication forks. Recently, scRepli-Seq has been developed to study replication timing in single cells(28, 29). This method relies on whole genome amplification followed by smoothing to identify replicated regions during S-phase (Figure 1F, Supplemental Figure 1H). In contrast, scEdU-seq relies on the identification of active DNA replication forks (Figure 1G). This allows for more accurate detection of sites where active DNA replication takes place, whereas scRepli-seq can only be used to determine which regions have already undergone DNA replication. Taken together, scEdU-seq allows high resolution profiling of DNA replication forks in single cells. Furthermore, these data allow accurate ordering of cells throughout S-phase.

**Fig. 1:**
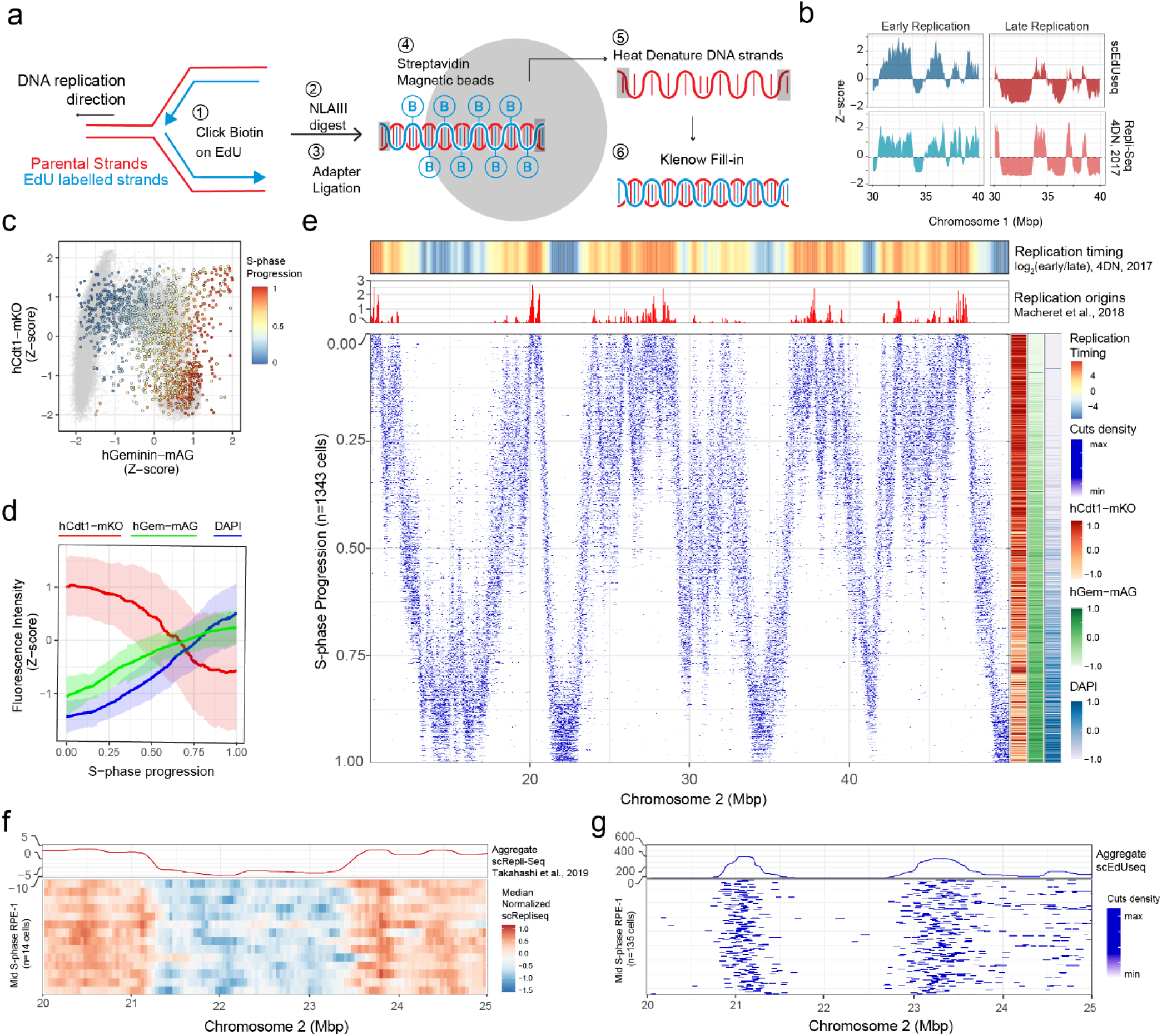
scEdUseq reveal ordered DNA replication profiles throughout S-phase. **a**. Representation of the scEdU-seq protocol. **b**. Z-scored genome-coverage tracks of early (*left*) and late (*right*) S-phase samples for both scEdU-seq (500 cell bulk, *top*) and Repli-Seq (400k cell bulk, *bottom*) treated with 120 min of EdU. **c**. Scatter plot showing FUCCI reporters pseudo colored by the S-phase progression based on scEdU-seq tracks and in gray the cell cycle distribution of cycling RPE-1 cells. **d**. Rolling mean of Z-scored fluorescence intensity of FUCCI reporters and DNA content (DAPI, *y-axis*) versus S-phase progression (*x-axis*) based on scEdU-seq tracks, ribbon indicates the standard deviation. **e**. Heatmap of scEdU-seq maximum normalized log counts for 1343 RPE-1 hTERT FUCCI cells ordered according to S-phase progression (*y-axis*) and binned per 400 kb bins (x-axis) for 50 megabase of chromosome 2. Top: heatmap showing log2-fold ratio of early to late Repli-Seq indicating replication timing (top) and bar graph showing replication origins of the same stretch of chromosome 2 (bottom). Right: scaled z-scored intensities of FUCCI reporters and DNA content (DAPI) ordered by S-phase progression. **f**. Heatmap of scRepli-Seq log2 median counts of middle S-phase cells binned per 40 kb (n = 14, *bottom*) and summed profile (*top*). **g**. Heatmap of scEdUseq maximum normalized log counts for middle S-phase cells binned per 5 kb (n = 135, bottom) and summed profile (top).

### Double pulse scEdU-seq allows DNA replication speed assessment

Next, we set out to estimate DNA replication speeds in single cells by using double-pulse EdU labeling (Figure 2A - top). After receiving two EdU pulses, the genome of a single cell in S-phase will be decorated with patches of EdU-containing DNA that are separated by a distance Δ*x*. The average replication velocity is then approximated by the distance Δ*x* divided by the inter-pulse duration Δ*t*. To systematically analyze this type of data, we make use of the pair correlation function(30), which reflects the distribution of all pairwise distances between EdU-containing reads in a single cell (Figure 2B - bottom). For two EdU pulses one would expect the pair correlation function to display two maxima: one at Δ*x* = 0 originating from pairs of reads that were labeled within one of the two pulses, and a second maxima is expected at 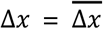 corresponding to read pairs that were labeled in separate pulses. It is the latter inter-pulse distances from which we can estimate the average length traveled by DNA replication forks in a single cell. Finally, we compute the average DNA replication fork speed for each cell by dividing the average distance traveled 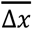 by the time between pulses Δ*t*. Interference from nearby active replication forks would not influence the value of 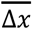. It could increase the overall background of the pair correlation function, because nearby replication forks will not be as regularly spaced as the double pulse signal. Consistently, the experimental pair correlation for a single cell exposed to a single EdU pulse only displays one maximum at Δ*x* = 0 (Figure 2B, Supplemental Figure 2). Single cells exposed to a double EdU pulse (i.e., Δ*t* = 45, 75 or 105 min) additionally show a second maximum. As expected, this second maximum shifts to larger values of Δ*x* as Δ*t* is increased (Figure 2C, top). The replication speed obtained from the experiments using different values of Δ*t* is comparable (Figure 2c, bottom). Subsampling the pairwise distances per chromosome shows similar distributions of 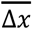 (Figure 2D). Taken together, we can use double pulse EdU labeling combined with pair correlation analysis to identify DNA replication speeds in single cells. To quantitatively model the pair correlation function we use a mixture model. This model separately fits distributions based on the intra-pulse differences (maximum at Δ*x* = 0), inter-pulse differences (maximum at 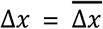) and the background (caused by, for example, nearby replication forks). The resulting distributions can subsequently be separated to infer the replication speed and the corresponding confidence interval in single cells(31) (Supplemental Figure 3 & 4).

**Fig. 2:**
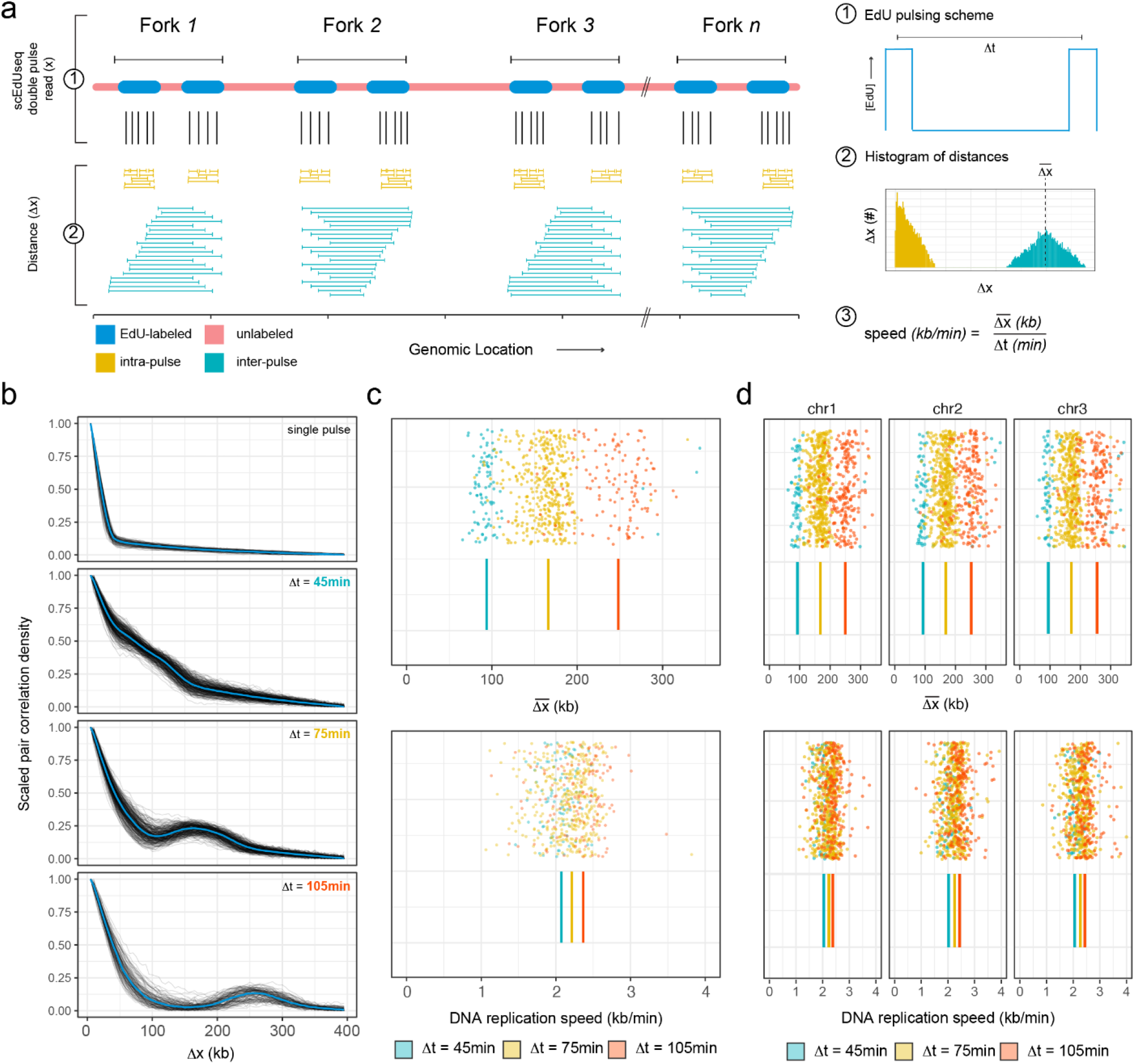
Double pulse scEdUseq allows DNA replication speed assessment. **a.** Schematic representation of the double pulse labeling scheme and subsequent analysis. b. Line plots of the pair correlations of single pulse, and double pulse labeling scheme with Δ t=45 (n=376), Δt=75 (n=347) and Δt=105 (n=149) minute labeling method. Every line is a single RPE-1 cell where the x-axis shows the binned distance and the y-axis the range-scaled density. Blue line indicates the mean density per bin. **c**. DNA replication distance estimates 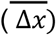 in kb for cells (Δt=45 (n=57), Δt=75 (n=299) and Δt=105 (n=103)) from Figure2b colored by labeling scheme. Ticks indicate the averages per labeling scheme (*top*). Distance estimates corrected for labeling resulting in DNA replication speeds kb/min (bottom). **d**. DNA replication speed estimates 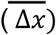 per chromosome (Δt=45 (n=57), Δt=75 (n=299) and Δt=105 (n=103)) from Figure2b colored by labeling scheme. Ticks indicate the average per labeling scheme (*top*). Distance estimates corrected for labeling scheme resulting in DNA replication speeds kb/min per chromosome (*bottom*)

### Transcription limits DNA replication speeds in early S-phase

We can approximate DNA replication speeds in single cells by using a double EdU pulse labeling strategy in combination with a mixture model. Several representative cells from a single-cell double EdU pulse experiment (Δt=75) demonstrate that the mixture model (Figure 3a, red dashed line) accurately describes the experimental pair correlation function (black solid line). Overall, we find DNA replication speeds in an expected range based on previous literature(32). Unexpectedly, we observe a large variability in replication speeds between individual cells (~1.5-fold difference, Figure 2c, d). A large part of this variability can be explained by the S-phase progression of single cells (Figure 3B). When analyzing DNA replication speeds during S-phase progression, we observe a steady average increase in DNA replication speeds suggesting an acceleration of replication as S-phase progresses (Figure 3B). DNA replication timing studies have shown that early replicating DNA is in proximity to actively transcribed regions of the genome(1). Since we observe the lowest average replication speeds in early S-phase, we hypothesize that these lower speeds might be correlated to transcription. Therefore, we use previously generated single-cell nascent RNA sequencing data on RPE-1 hTERT cells to test this assumption (Supplemental Figure 5A-D)(33). We observe the highest levels of nascent transcription in regions of the genome that are covered by scEdU-seq domains at the start of S-phase (Figure 3C). In addition, both the number of transcribed regions as well as transcription levels decrease as DNA replication progresses over S-phase. The presence of high levels of transcription in early S-phase therefore correlates to lower DNA replication speeds (Figure 3B,C). This could imply that transcription limits DNA replication speeds. If active RNA polymerase 2 (RNAP2) transcription reduces DNA replication speeds, we would expect an increase in DNA replication speeds in early S-phase by inhibiting transcription. In order to assess DNA replication speeds without active RNAP2 transcription, we treat cells with the transcription inhibitor 5,6-Dichlorobenzimidazole 1-β-D-ribofuranoside (DRB, 60 min, Supplemental Figure 5E-I) between the two EdU pulses, which does not alter either initiation zones or replication timing (Supplemental Figure 5J-K). We observe an overall increase in DNA replication speeds in RNAP2 inhibited cells versus DMSO treated cells (Figure 3D, bottom). Furthermore, this difference in speed is a result of increasing DNA replication speeds during early S-phase (Figure 3D, top). In summary, DNA replication accelerates over S-phase, which is, in part, a result of limiting DNA replication speeds during early S-phase by RNAP2 transcription.

**Fig. 3:**
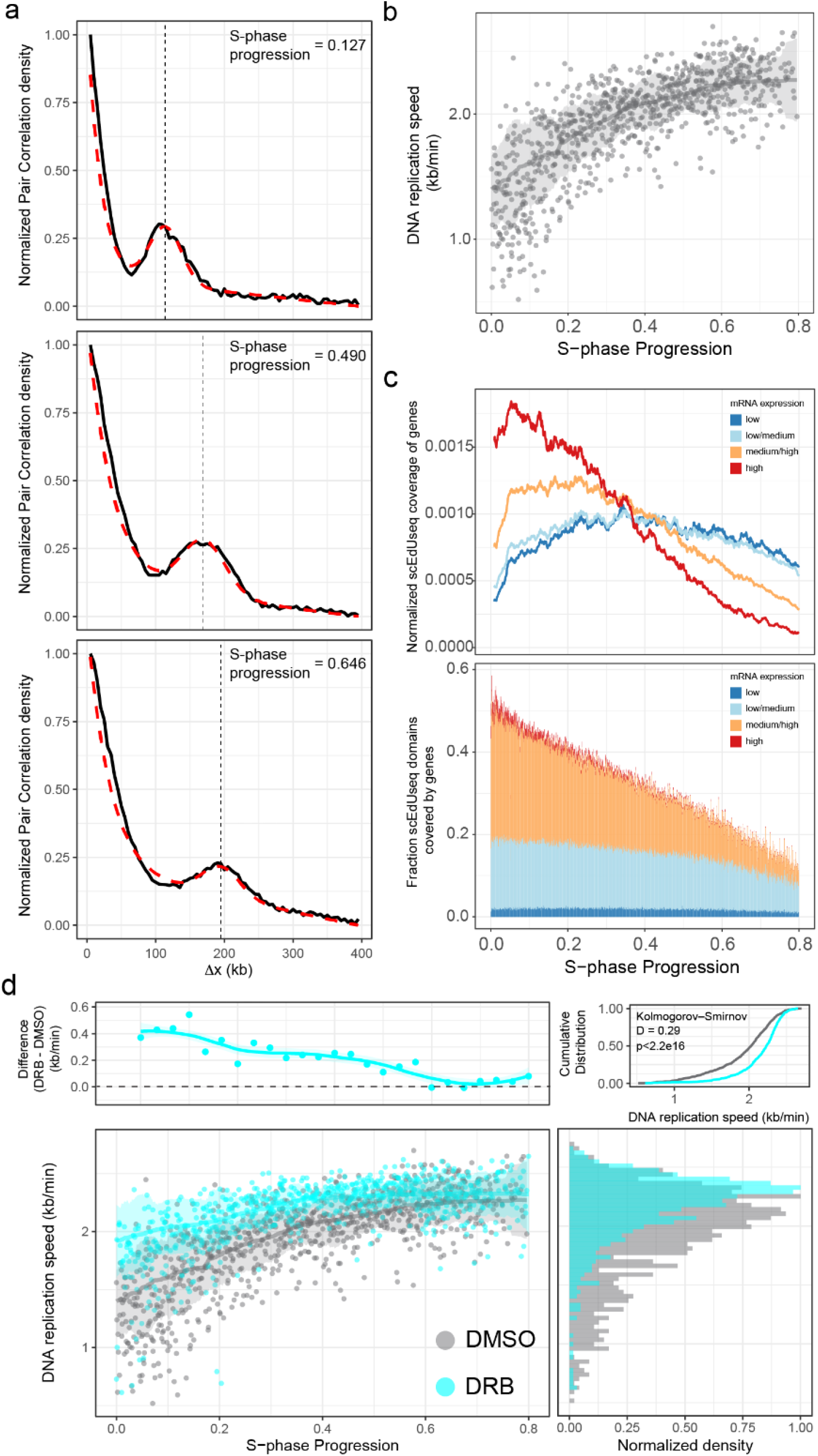
Transcription limits DNA replication speeds in early S-phase. a. Single-cell pair correlations (black line) with fitted model (dashed red line) of representative early-(top), middle-(middle) and late S-phase (bottom) RPE-1 cells labeled with the Δt=75min scheme. The x-axis shows the binned distance in kilobase (kb) and the y-axis the range-scaled density. b. DNA replication speed over S-phase in RPE-1 (n=326) treated with DMSO subjected to Δt=75 labeling scheme. Speed estimates in kb/min (y-axis) over S-phase progression (x-axis). Every dot is a cell, the line indicates a rolling-window median smooth and the ribbon the standard deviation around the median. c. Nascent RNA sequencing from S-phase RPE-1 cells. Rolling window smoothened normalized scEdU-seq coverage of genes (y-axis) over S-phase progression (*x-axis*) colored by expression level (*top*). Fraction of scEdU-seq domains covered by expressed genes (*y-axis*) over S-phase progression (*x-axis*) colored by expression level (stacked). d. DNA replication speed over S-phase in RPE-1 treated with DMSO (gray, n=326) or DRB (cyan, n=713) subjected to Δ t=75 labeling scheme (*bottom-left*). Difference in DNA replication speeds between DMSO and DRB in kb/min (*y-axis*) over S-phase progression (*x-axis, top-left*), marginal density (*x-axis*) of DNA replication speed in kb/min (*y-axis*) colored for DMSO-(*gray*) or DRB-treated cells (*cyan, bottom-right*) and cumulative distribution of marginal speed density (*top-right*).

### Transcription-coupled damage decreases DNA replication speeds

RNAP2 activity has been correlated to a variety of DNA damage, for example, by generating SSBs through Topoisomerase I cleavage complexes or repair of bulky adducts by transcription-coupled nucleotide excision repair(34, 35). Moreover, conflicts between the DNA replication fork and transcription machinery results in the formation of RNA:DNA hybrids, which result in double-strand breaks if improperly handled(36). Indeed, short inhibition of RNAP2 (1 hr) during S-phase results in a reduction of DNA damage as assayed by flow cytometry (Figure 4A, γH2AX). This suggests that RNAP2 activity, at least in part, causes transcription-coupled DNA damage during S-phase. The activity of the DNA damage sensor PARP-1 is stimulated by a wide variety of DNA damage lesions(37). Since PARP-1 activity has previously been linked to DNA replication speeds(32), we reason that the decrease of DNA replication speed in early S-phase might be due to transcription-coupled DNA damage and subsequent PARP activation. We observe a decrease in the level of pan ADP-ribose, the modification deposited by PARP enzymes (Figure 4A, pan ADP-ribose). This suggests that RNAP2 transcription not only induces DNA damage, but also activates PARP. In order to explore how transcription-coupled DNA damage might affect DNA replication speed in single cells, we make use of the PARP inhibitor, Olaparib. First, we treat wild-type RPE-1 cells with a PARP inhibitor and observe very similar behavior compared to RNAP2 inhibition (Figure 4B, Supplemental Figure 6A-E). Overall DNA replication speeds are higher in PARP-inhibited cells without altering either IZ or replication timing. Additionally, the most notable difference of DNA replication speeds occurs in early S-phase suggesting a connection to RNAP2 transcription. To directly address the role of PARP activity in regulating DNA replication speeds, we set out to hyperactivate PARP-1 by generating an RPE-1 cell line in which the gene XRCC1 was knocked out (XRCC1Δ RPE-1, Supplemental Figure 6 G,H). The XRCC1 protein is required for efficient repair of DNA damage. In the absence of this protein, steady-state levels of DNA damage increases causing PARP hyperactivation, which eventually leads to cerebral ataxia(38) (Figure 4C-E). XRCC1Δ RPE-1 cells have a lower proportion of EdU+ cells compared to wild-type cells, which is partially mitigated by the addition of a PARP inhibitor (Figure 4F). This implies that excessive PARP signaling in XRCC1Δ RPE-1 results in lower DNA replication speeds. In line with this observation, the XRCC1Δ RPE-1 cells display overall lower DNA replication speeds compared to wild-type cells (Figure 4G, Supplemental Figure 6I,J). In contrast to RNAP2 or PARP inhibition, we observe a ubiquitous decrease in DNA replication speeds in all cells and not just early S-phase cells. This suggests that hyperactivation of PARP, outside of transcribed regions, results in lower DNA replication speeds. In addition, we can rescue the global decrease of DNA replication speeds in XRCC1Δ RPE-1 by the addition of a PARP inhibitor (Figure 4H). This suggests that PARP activity is critical in regulating DNA replication speeds in response to transcription-coupled DNA damage.

**Figure 4:**
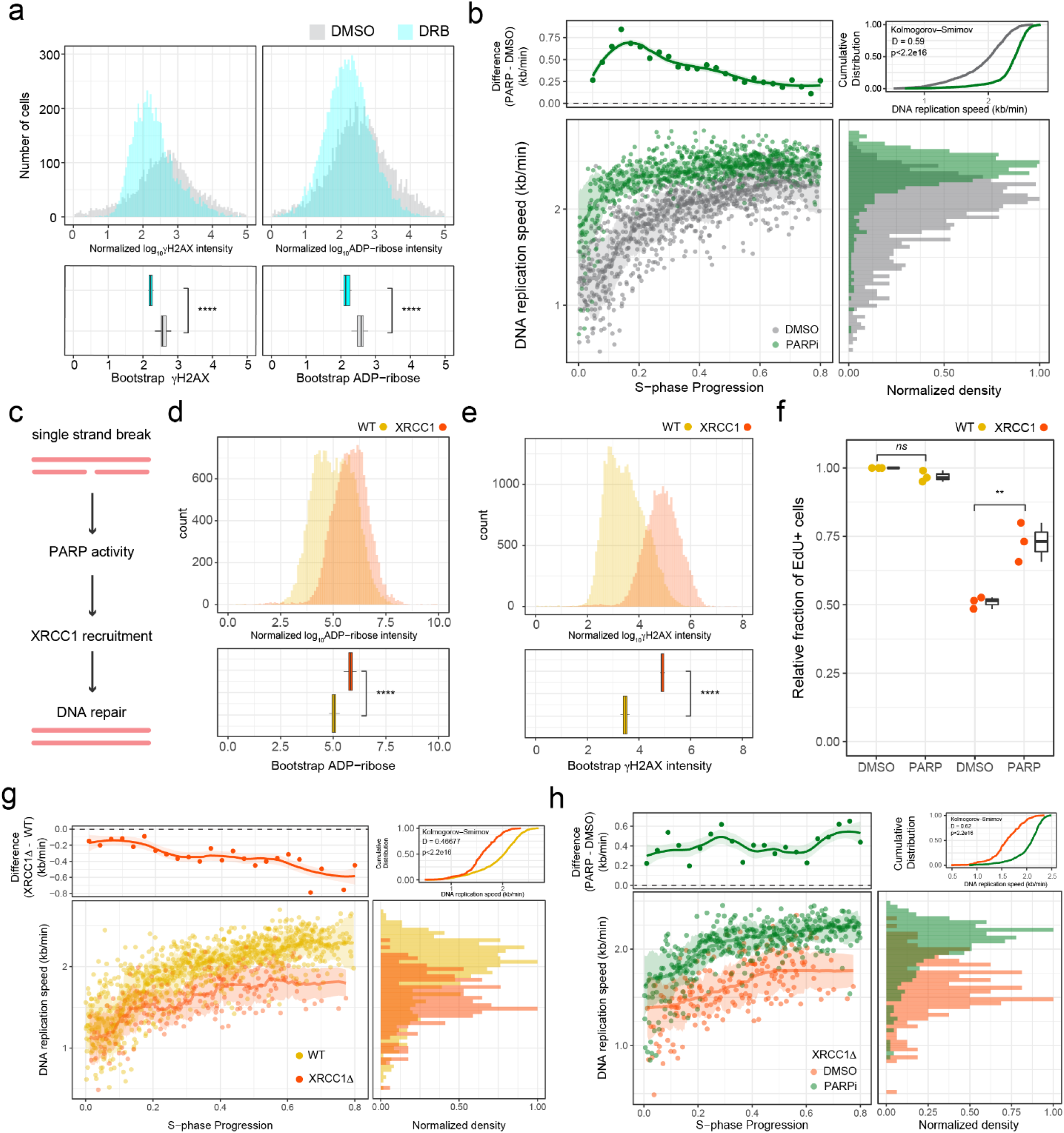
Transcription-coupled damage decreases DNA replication speeds. a. Density (y-axis) of scaled z-score log10-transformed ADP-ribose (left) or yH2AX (right) intensity (x-axis) for either DMSO (gray) or DRB-treated cells (cyan, top) and bootstrapped mean of the densities in the top of the panel (bottom). b. DNA replication speed over S-phase in RPE-1 treated with DMSO (gray,n=326) or 24hr PARPi (green, n=766) subjected to Δt=75 labeling scheme (*bottom-left*). Difference in DNA replication speeds between DMSO and PARP in kb/min (*y-axis*) over S-phase progression (*x-axis, top-left*), marginal density (*x-axis*) of DNA replication speeds in kb/min (*y-axis*) colored for DMSO-(*gray*) or PARP-treated cells (*cyan, bottom-right*) and cumulative distribution of marginal speed density (*top-right*). c. Schematic representation of DNA damage signaling and repair of single strand breaks (SSB). d,e. Density (y-axis) of scaled z-score log10-transformed ADP-ribose (left) or γH2AX (right) intensity (x-axis) for either WT (yellow) or XRCC1Δ RPE-1 cells (orange, top) and bootstrapped mean of the densities in the top of the panel (bottom). f. Relative fraction of EdU positive cells (y-axis) for DMSO vs 4hr PARPi-treated cells (x-axis) colored by WT (yellow) or XRCC1Δ RPE-1 cells (red). g. DNA replication speed over S-phase in RPE-1 WT (yellow, n=326) or XRCC1Δ RPE-1 (red, n=187) subjected to Δt=75 labeling scheme (*bottom-left*). Difference in DNA replication speeds between WT RPE-1 and XRCC1Δ RPE-1 in kb/min (*y-axis*) over S-phase progression (*x-axis, top-left*), marginal density (*x-axis*) of DNA replication speeds in kb/min (*y-axis*) colored for WT RPE-1 (*yellow*) or XRCC1Δ RPE-1 (*red, bottom-right*) and cumulative distribution of marginal speed density (*top-right*). h. DNA replication speed over S-phase in XRCC1Δ RPE-1 treated with DMSO (red, n=187) or PARP (green, n=393) subjected to Δt=75 labeling scheme (*bottom-left*). Difference in DNA replication speeds between DMSO and 4hr PARPi in kb/min (*y-axis*) over S-phase progression (*x-axis, top-left*), marginal density (*x-axis*) of DNA replication speeds in kb/min (*y-axis*) colored for DMSO-(*red*) or PARP-treated cells (*green, bottom-right*) and cumulative distribution of marginal speed density (*top-right*)

## Discussion

Taken together, we developed a method with which we can profile DNA replication forks and their speeds in single cells. We show that DNA replication speeds accelerate during S-phase. Reduced DNA replication speeds at the start of S-phase occur in genomic regions with high levels of RNAP2 transcription. Inhibition of RNAP2 transcription increases DNA replication speeds at these locations. We find that inhibition of RNAP2 results in both lower PARP activity as well as lower DNA damage. We continue to show that lowering PARP activity allows for higher DNA replication speeds, specifically in early S-phase. In addition, the hyperactivation of PARP in RPE-1 cells lacking XRCC1 results in a genome-wide decrease of DNA replication speeds. We can revert this decrease by lowering PARP activity, indicating a direct role in regulating DNA replication speeds. Overall, this implies that transcription-coupled DNA damage increases PARP activity, which in turn reduces DNA replication speed.

Our data suggests crosstalk between DNA replication fork speeds and transcription through the activity of PARP enzymes. Interestingly, proteomic profiling has identified DNA replication and transcription as the two biological processes regulated through ADP-ribosylation by PARP enzymes(39). Several core DNA replication factors have been implicated in regulating fork speeds(32, 40, 41). However, which specific components are regulated by ADP-ribose to reduce replication fork speeds remains to be uncovered. scEdUseq in combination with a small scale genetic perturbation screen would be uniquely suited to identify these components. In addition, several studies have shown the role of PARP activity in directing transcriptional activity(39, 42). As of yet, we are unable to identify the direct effect of transcription on DNA replication. Conversely, we can not assess the effect of active DNA replication on RNAP2 transcription. Therefore, integration of transcriptome profiling with scEdU-seq would allow the coupling of RNAP2 transcription dynamics to the proximity of active DNA replication. Furthermore, we would be able to discern dynamic or transient cell states by transcriptome profiling and relate these to rare DNA replication phenotypes (e.g. fast DNA replication speeds in early S-phase).

Potential limitations of related techniques, including scEdU-seq, is that they rely on incorporation of non-natural nucleotides which might disturb the system in non-biologically relevant ways. Searching for endogenous read-outs for active DNA replication in single cells would be important to circumvent such a potential limitation.

We expect that scEdU-seq will have applications to identify single-cell DNA replication timing as well as replication speeds in a wide range of biological systems. In addition, unscheduled DNA replication such as homologous recombination as well as replication intermediates from previous cell cycles could potentially be detected through scEdU-seq. Finally, extracting transcriptome profiles in conjunction with DNA replication profiles will allow uncovering of molecular crosstalk as well as identification and characterization of rare events.

## Methods

### Cell lines and Reagents

RPE-1 hTERT FUCCI and RPE-1 hTERT XRCC1Δ cells were cultured in DMEM/F12 supplemented with 10% FBS (Gibco), 1 × GlutaMAX (Gibco) and 1 × Pen-Strep (Gibco) at 37 °C with 5% CO2. RPE-1 cells routinely tested negative for Mycoplasma contamination and were not authenticated. Counting of cells was performed with the Bio-Rad TC-20 Cell Counter. The following chemicals were used: EU (100 μM, Invitrogen), EdU (10 μM, Invitrogen), Olaparib (AZD2281, 10 μM, Cell Signaling Technology), DRB (10 μM, Sigma), and DAPI (10 mg/mL, Thermo-Fisher) and SN-38 (used at indicated concentrations, SelleckChem)

### XRCC1Δ knockout generation in RPE-1 hTERT cells

The guideRNA was designed using CRISPOR (crispor.tefor.net) against the 2nd exon of XRCC1. The primers 5’-CACCGAGACACTTACCGAAAATGGC-3’ and 5’-AAACGCCATTTTCGGTAAGTGTCTC-3’ (Integrated DNA Technologies,Belgium). were cloned into pX330(43). Cells were co-transfected with pDonor-Blast(44) and the pX330-XRCC1. Cells were allowed to recover for 72 hours and selected with Blasticidin. Clones were picked and expanded. The picked clones were validated by Western Blot analysis as previously described(45) and probed with CDK4 (Santa-Cruz Biotechnologies, sc-260, 1:1000) and XRCC1 (Abcam, ab1838, 1:1000) primary antibodies.

### SN-38 Proliferation assay

500 RPE-1 hTERT or RPE-1 hTERT XRCC1Δ were plated in a MW-96 and treated with increasing concentrations of SN-38 for 120 hours. Cell viability was measured at the end of the experiment with CellTiter-Glo® Cell Viability Assay according to the manufacturer’s protocol

### Single and Double pulse EdU treatment

For single pulse experiments, 1.5 × 10^6^ cells were treated with 10 μM EdU for 15 minutes, trypsinized and washed with 1×PBS, followed by fixation in 70% ice-cold ethanol. Double pulse experiments were performed by treating 1.5 × 10^6^ cells with 10 μM EdU for 15 minutes, subsequently cells were washed 3 times with DMEM/F12 medium allowed to recover for indicated time periods (e.g. 30, 60 or 90 minutes). Finally, cells were treated with a second pulse EdU (10 μM), trypsinized, washed in PBS and fixed in 4 mL 70% ethanol.

Cells were stored at −20°C for up to 24 hours. For longer storage periods up to 3 months, cells were stored in 4 mL Storage Buffer (150 mM NaCl, 20 mM HEPES, 2 mM EDTA, 25 mM Spermidine with 10% DMSO) at −20°C.

### Azide-PEG3-Biotin EdU Click reaction

Eppendorf Protein lo-bind 0.5 mL tubes were precoated with 0.25% BSA in PBS. Afterwards 500 mL of cells, in either 70% Ethanol or storage buffer, were pelleted for 3 min at 600g. The cells were resuspended in 0.25% BSA in PBS and left to block for 30 minutes at 4°C. Following blocking, cells were pelleted and the click reaction was performed *in situ* with 50 uL reactions with the EdU Click 647 Imaging Kit (brand) according to manufacturer’s protocol with some alterations. The Azide-647 was replaced with Azide-PEG3-Biotin (Sigma, 10 mM) and supplements with 6 mM THTPA (Jena Bioscience).

### FACS

Following click reaction, RPE-1 cells were washed once in 1× PBS0, resuspended in PBS0 with 0.25% BSA (Thermo Fisher) and 10 μg/mL DAPI, and passed through a 20-μm mesh. Single cells were index sorted using a BD FACS Influx with the following settings: sort objective single cells, a drop envelope of 1.0 drop, a phase mask of 10/16, extra coincidence bits of maximum 16, drop frequency of 38 kHz, a nozzle of 100 μm with 18 PSI and a flow rate of approximately 100 events per second, which results in a minimum sorting time of approximately 5 min per plate.

Doublets, debris, and dead cells were excluded by using the forward and side scatter and DAPI channel. For the hTERT RPE-1 FUCCI cells, the measurements in the DAPI channel were used to enrich S-phase cells. The intensities in the monomeric Azami-Green (mAG) and monomeric Kusabira-Orange 2 (mKO2) as well as DAPI channels were acquired and later used for data analysis. Single cells were sorted into 384-well hardshell plates (Bio-Rad) containing 5 μl of light mineral oil (Sigma-Aldrich).

### Library Preparation

Library construction progressed through three general steps (Figure 1a): cell lysis and NLAIII digestion, End Repair and A-tailing followed by Adapter Ligation, and pooling and purification. Reagents were dispensed to 384 microwell plates using either the Nanodrop II (Innovadyne Technologies) or the Mosquito (TTP Labtech). Plates were spun at 2,000g for 2 minutes after each liquid transfer step.

### Cell Lysis and NLAIII digestion

After sorting, single cells were lysed in 100 nL lysis mix (10 nL 1X Cutsmart buffer (NEB), 10 nL Proteinase K (Ambion), 80 nL H_2_O). Plates were spun at 2000g for 2 min and incubated for 2 hours at 55°C and the Proteinase K was heat inactivated for 20 minutes at 80°C. The genome was digested with 100nL NLAIII mix (10 nL 1X Cutsmart buffer (NEB), 10 nL NLAIII (Ambion), 80 nL nuclease-free H_2_O) at 37°C for 4 hours and heat inactivated for 30 minutes at 65°C.

### End Repair and A-tailing followed by Adapter Ligation

In order to end repair NLAIII overhang, we incubated single cell with 100 nL of end repair mix (1.6 nL Klenow Large Fragment (NEB), 1.6 nL T4 PNK (NEB), 4 nL 10 mM dNTPs, 2.3 nL 100 mM ATP, 6.6 nL 25 mM MgCl2, 5 nL PEG8000 (50%, NEB), 1.2 nL BSA 20 ng/ml (NEB), 23.3 nL 10X PNK buffer (NEB) and 54.2 nL nuclease-free H_2_O) for 30 minutes at 37°C and heat inactivated both enzymes for 20 minutes at 75°C. In order to ligate adapters with a T-overhang, we A-tailed the end-repaired genomic DNA fragments with 100 nL of A-tailing mix (0.66 nL AmpliTaq 360 (Thermo-Fisher), 0.66 nL 100 mM dATP, 16.6 nL 1M KCl, 5 nL PEG8000 (50%, NEB) 0.5 nL BSA (20ng/ml, NEB) and 77.2 nL nuclease-free H_2_O) for 15 minutes at 72°C. Finally, A-tailed fragments were ligated to 50 nL of 5 mM T7 promoter containing adapters(21) with cell barcodes and UMI with 150 nL ligation mix (25 nL T4 DNA ligase (400.000U, NEB), 3.5 nL MgCl_2_, 10.5 nL TRIS pH 7.5 (1 M, Gibco), 5.25 nL DTT (1M, Thermo-Fisher), 3.5 nL ATP (100 mM, Thermo-Fisher), 10 nL PEG8000 (50%, NEB), 1 nL BSA (20 ng/ml NEB) and 91.25 nL nuclease-free H_2_O) for 20 minutes at 4°C followed by 16 hours at 16°C and heat inactivated for 20 minutes at 65°C.

### Pooling and purification of EdU fragments

The contents of each plate were collected in mineral oil pre-coated VBLOK200 reservoirs (ClickBio) by centrifuging at 300g for 1 min. The aqueous phase (approximately 200 mL per plate) was collected and separated from residual mineral oil by centrifugation. EdU-PEG3-Biotin containing DNA molecules were affinity purified by Streptavidin MyOne C1 Magnetic Beads (Invitrogen) according to the manufacturer’s protocol. Subsequently, we retrieved the complementary strand of the EdU-PEG3-Biotin containing DNA strand by heat denaturation at 95°C. While ramping down the temperature (0.1°C/s) to 20°C, we annealed an oligo (5’-ATGCCGGTAATACGACTCAC-3’) complimentary to the constant adapter sequence region in oligo annealing buffer (20 mM TRIS pH 8, 1mM MgCl_2_, 100mM NaCl). Next, we extended the primer to generate double-strand DNA with Klenow Large fragment mix (1X NEB Buffer 2, 50 mM dNTPs, 0.5U Klenow Large Fragment) for 45 minutes at 25°C and heat inactivated for 20 minutes at 75°C. DNA fragments were purified with Ampure XP beads (Beckman Coulter) at a sample to beads ratio 1:1 and resuspended in 7 mL nuclease-free H_2_O.

### Library amplification by IVT and PCR

Pre-amplified libraries were linearly amplified by MEGAscript™T7 Transcription Kit (Thermo-Fisher) for 12 hours at 37°C. Template DNA was removed by the addition of 2 μl TurboDNAse (Thermo-Fisher) for 15 min at 37°C. Amplified RNA (aRNA) was fragmented for 2 min at 94°C with Fragmentation Buffer (5X concentrated; 200 mM Tris-acetate pH 8.1, 500 mM KOAc, 150 mM MgOAc). aRNA was directly cooled back down to 4°C on ice and 50 mM EDTA was added to stop fragmentation. The fragmented RNA is purified using RNA Clean XP beads (Beckman Coulter) at 1:1 beads to sample ratio and eluted in 12 mL of nuclease-free H_2_O. Next, 5 mL aRNA was converted to cDNA by reverse transcription in two steps. First, the RNA is primed for reverse transcription by adding 0.5 μl dNTPs (10 mM) and 1 μl random hexamer RT primer 20 μM (5’-GCCTTGGCACCCGAGAATTCCANNNNNN-3’) at 65°C for 5 minutes followed by direct cooling on ice. Second, reverse transcription is performed by addition of 2 μl First Strand buffer, 1 μl DTT 0.1 M, 0.5 μl RNAseOUT and 0.5 μl SuperscriptII and incubating the mixture at 25°C for 10minutes followed by 60 minutes at 42°C and 20 minutes at 70°C. Single stranded cDNA was purified from aRNA through incubation with 0.5 μl RNAseA (Thermo-Fisher) for 30 min at 37°C. Finally, cDNA was amplified by PCR, which also attaches the Illumina small RNA barcodes and handles, by adding 25 μl of NEBNext Ultra II Q5 Master Mix (NEB), 11 μl Nuclease free water and 2 μl of RP1 and RPIx primers (10 μM). DNA fragments were purified, twice, with Ampure XP beads (Beckman Coulter) at a sample to beads ratio 0.8:1 and resuspended in 10 mL nuclease-free H_2_O. Abundance and quality of the final library are assessed by Qubit and Bioanalyzer.

### Sequencing

Libraries were sequenced using v2.5 chemistry on a NextSeq 500 or NextSeq2000 (Illumina; NextSeq Control Software version 2.2.0.4; RTA version 2.4.11) with 100 cycles for Read 1 (cell index and UMI) and and 100 cycles for Read2 (sample index).

### Flow Cytometry

Apoptosis analysis was performed by using Annexin-V-APC kit (BioLegend) as described in manufacturers protocol for all treatment conditions and cell lines. Cell cycle analysis was performed by fixing RPE-1 hTERT FUCCI or RPE-1 hTERT XRCC1Δ cells in 70% ethanol and counterstaining with DAPI (10 μg/mL, 20min on ice). For nascent RNA labeling, we treated cells with EU (200 μM) for 1 hour and fixed the cells in 70% ethanol. For DNA replication labeling, we incubate cells for 30 minutes with 10 μM EdU. For both EU and EdU labeling, we used the EdU Click 647 Imaging Kit according to the manufacturer’s protocol. For γH2AX staining, we used the FITC-conjugated γH2AX (1:500, Millipore). panADP-ribose binding reagent (1:1500, Sigma-Aldirch, MABE1016) was used in combination with Donkey anti-Rabbit IgG Secondary Antibody with different Alexa Fluor conjugation depending on experimental condition and cell line used.

### Read processing

Processing of raw fastq to count tables was performed by SingleCellMultiOmics v0.1.25 (https://github.com/BuysDB/SingleCellMultiOmics).

First, fastq files were demultiplexed, which adds UMI, cell, sample and sequencing indices to the header of the fastq. Cell barcodes and UMIs with a hamming distance of 1 were collapsed. Next, the adapter sequences were trimmed from each read with cutadapt. Subsequently, reads were mapped with bwa (BWA MEM algorithm) to Ensembl release 97, GRCh38.p12 for *Homo Sapiens*, the bam outputs were sorted with samtools. Mapped reads were subjected to molecule assignment, which generates tags for NLAIII restriction site position and integrates cell barcode, UMI, library, strand and genomic position of NLAIII restriction site into one tag. This integrated molecule tag allows for deduplication of reads and generation of long format tables. These tables were filtered on the presence of a NLAIII restriction site, a mapping quality >30, the molecule has a pair of reads assigned, the molecule is unique, and should not have alternative alignment positions in the genome.

### Single-cell DNA replication analyses and plotting

All data analysis was done in R using the *tidyverse* and *data.table* packages unless otherwise mentioned.

### Comparison between Repli-Seq and scEdU-seq

Bulk scEdU-seq samples were generated by collecting 500 early or late S-phase RPE-1 hTERT FUCCI cells treated with EdU for 120 minutes. Cells were processed similarly as scEdU-seq libraries. We retrieved bulk Repli-Seq for RPE-1 hTERT cells from the 4D Nucleome(27), which was generated by the Gilbert lab. Subsequently, the samples were binned with a 50 kb resolution and reads per bin were Z-scored per sample. Z-scores were used for comparative plotting of traces as well as computing Spearman Correlation between samples.

### DNA replication origin data from RPE-1 hTERT

Raw DNA replication origin data (EdUseq-HU) for RPE-1 hTERT was used from Macheret et al(26) (BioProject PRJNA397123). Raw FASTQ files were trimmed and mapped to Ensembl release 97, GRCh38.p12 with bwa (mem algorithm). The data were binned with a 50 kb resolution and reads per bin were Z-scored, which were subsequently plotted over selected genomic regions.

### scEdU-seq and scRepli-Seq comparison

Processed single-cell Repli-Seq for Mid S-phase RPE-1 hTERT cells was acquired from the Gene Expression Omnibus (GSE108556). Mid S-phase cells for scEdUseq were selected on S-phase progression (0.37;0.47). Binned single cell tracks for both methods were plotted for a region of chromosome 2 (40kb for scEdUseq and 200kb for scRepliSeq). In addition, aggregate tracks of scRepli-Seq and scEdUseq were generated by the addition of values per bin and plotted above single-cell tracks.

### S-phase ordering

First, cells were filtered by the average counts per 100kb bin with a lower threshold (single pulse 0.37; double pulse 0.08) and an upper threshold (single pulse 2.72; double pulse 12.18), and deviance of poisson behavior defined as

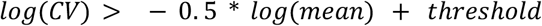

where the threshold was set to 0.1. An exponential mixture model was fitted on the distances between successive reads for each single cell separately using the R-package *flexmix(31)*. Subsequently, reads with a posterior probability of higher than 0.5 for the distances to their first neighbors, were used for S-Phase ordering. Next, we performed a gaussian kernel smoothing (sd of 8333.333) for the remaining reads, after which the pairwise overlap coefficient was calculated between all cells on a *per* chromosome basis. The overlap coefficient was converted to a distance as follows:

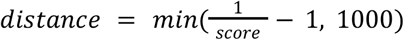

and averaged over chromosomes. The resulting distance was embedded in 1 dimension using Uniform Manifold Approximation and Projection (UMAP) implemented by the R-package *umap*. This UMAP computation was repeated 100 times and the resulting UMAP axis was converted to a Z-score. To prevent a flipping of the direction S-phase progression the runs that had an average Spearman rank correlation lower than 0.85 with the other runs were discarded. Finally, to determine if a cell could be placed on the ordering definitively, placings were considered clustered if the smallest distance between two successive placings was smaller than 0.1, and cells were kept if the biggest cluster contained at least 80% of the successful runs.

### Pair correlation

The pair correlation was calculated as the pairwise distances between all reads in one cell per chromosome. To display the count was calculated per 5kb bin, distances bigger than 400kb (except FigureS 3 for which the max was 1Mb) were discarded and the total counts were either sum normalized or range-scaled between 0 and 1.

### Mixture model fits

A mixture model with 4 (uniform, exponential, halve-normal and normal distribution) components was fitted per cell using a custom EM-algorithm with soft labels written in C++ and implemented using the R-package *Rcpp(46)*. In the Maximization-step the parameters of the component distributions were updated using the weighted mean for the exponential and normal distribution and the weighted variance for the halve-normal and normal distribution. Additionally, the mean of the exponential component was restricted to be above 1000, and the prior probability of the exponential component was restricted to be above 0.01. The algorithm was run until a relative tolerance of 10^−8^ or a maximum of 100 iterations.

### Pair correlation simulation

In the case of sampling with equal intensity, to simulate the pair correlation first the read locations were drawn from a uniform distribution with a minimum of −200 and a maximum of 200, where the number of reads was drawn from a poisson distribution with a mean of 80. Then reads *r* were kept with the following logic:

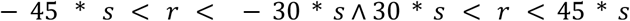

where the scaling *s* of the window was drawn from a truncated (at 0) normal distribution with a given mean and variance.

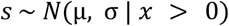

In the case of unequal sampling intensity first the double window was defined and scaled with a speed factor drawn from a (truncated at 0) normal distribution with a given mean and variance.

Then the read locations were drawn from a uniform distribution where the minimum and maximum were scaled according to the speed factor, and the number of reads was again drawn from a poisson distribution with a mean of 80. Reads falling outside the earlier defined window were again discarded.

For both scenarios this was done 2000 times for every combination of mean and variance of the ground truth speed distribution, after which the pair correlation was calculated as described before.

Subsequently, fitted speed estimates were corrected for poisson sampling artifacts by fitting a loess smoothing surface to the simulated data. After which, we predict the input mean or standard deviation parameter using the output mean and standard deviation Finally, this model predicts the ground truth mean and standard deviation from the measured experimental mean and standard deviation.

#### HMM pulse segmentation

To determine the part of the genome that is undergoing replication in a single pulse experiment, a 2-state Hidden Markov Model was used to segment the genome into foreground and background. We used the R-package *mhsmm(47)* to fit a Hidden Semi Markov Model with exponential emission distributions and a gamma sojourn distribution per cell on the distances between neighboring reads, where each of the chromosomes were used as separate observations. Then to get the most likely sequence of states the viterbi algorithm implemented in the mhsmm package was used.

#### scRNA-seq data analysis

Count tables from Battich et al.(33) were filtered for the EU-labeled fraction of mRNA, and cells were assigned to S-phase if the cell cycle progression score (see Battich et al.) was over 0.333 or below 0.75. The total counts per gene for the S-phase pseudo-bulk were then transformed by adding 1, log10 transformed and rounded.

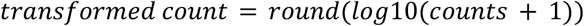

The genomic location was added to the genes using the hg38 ensembl release 106. In the case of overlapping genes the one with the higher count was given priority for the overlapping portion. The overlap of the segmented single-pulse data with the expressed genes was calculated by first identifying overlaps using the overlap function from the data.table package and then calculating the overlap.

## Acknowledgements

We thank R. van der Linden for assistance during experiments. We thank P. Knipscheer, J. Garaycoechea and F. Mattiroli, for providing valuable feedback on the manuscript. We also thank all members of the van Oudenaarden lab for scientific discussions. This work was supported by a European Research Council Advanced grant (ERC-AdG 742225-IntScOmics) and Nederlandse Organisatie voor Wetenschappelijk Onderzoek (NWO) TOP award (NWO-CW 714.016.001). This work is part of the Oncode Institute, which is partly financed by the Dutch Cancer Society. In addition, we thank the Hubrecht Sorting Facility and the Utrecht Sequencing Facility, subsidized by the University Medical Center Utrecht, the Hubrecht Institute, Utrecht University and The Netherlands X-omics Initiative (NWO project 184.034.019).

## Contributions

J.v.d.B. and A.v.O. conceived and designed the project. J.v.d.B. developed the experimental scEdUseq protocol and performed experiments. V.v.B. developed the statistical and analytical framework to analyze scEdUseq data with the help of J.v.d.B. and A.v.O. J.v.d.B. V.vB. and A.v.O analyzed the data. J.v.d.B., V.v.B. and A.v.O. discussed and interpreted results. J.v.d.B. wrote the manuscript with feedback from A.v.O. and V.v.B.

## Data availability

Raw sequencing data, metadata and count tables have been made available in the Gene Expression Omnibus under the accession number GSE211037. Data for comparisons to scEUseq and scRepli-Seq were downloaded from Gene Expression Omnibus accessions GSE128365 and GSE108556. Raw sequencing data of DNA replication origins was downloaded from SRA (PRJNA397123). Data for Replication timing was downloaded from the 4D nucleome project (4DNBSKYMY5XL)

## Code availability

All scripts to process raw data and generate figures are available at https://github.com/vincentvbatenburg/scEdU-seq

## Supplemental Figures

**Supplemental Figure 1:**
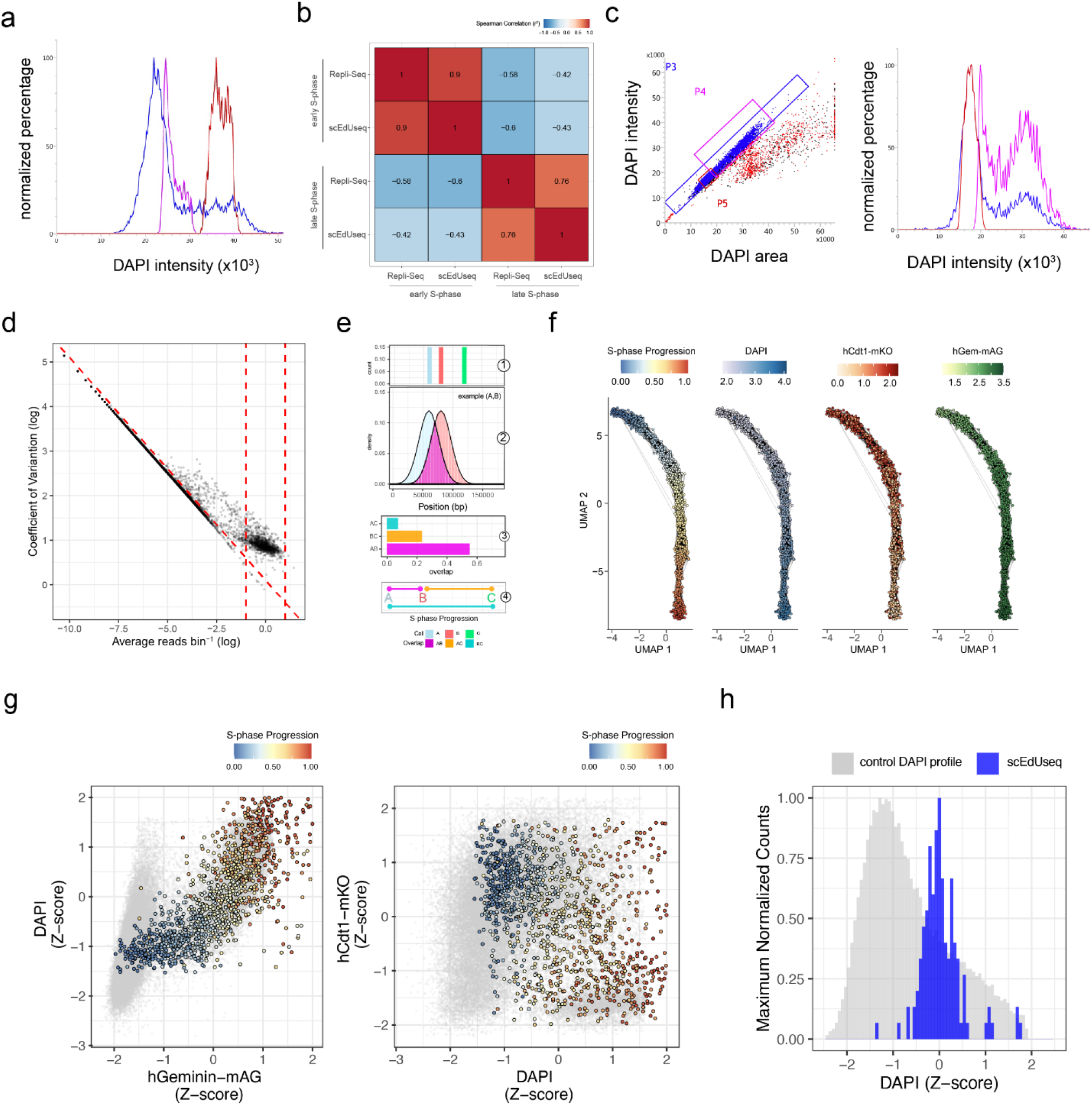
Supplemental Figure to Figure 1. **a**. Enrichment gates for DNA content (DAPI) to sort early (*purple*) and late (*brown*) superimposed over the cell cycle distribution of cycling RPE-1 cells (*blue*). **b**. Spearman rank correlation heatmap comparing early vs late S-phase sorted samples between Repli-Seq (400.000 cells) and scEdU-seq (500 cells) with 120 minutes EdU treatment. **c**. Enrichment gate for DNA content (DAPI) to single cells (P4, purple) superimposed over the cell cycle distribution of cycling RPE-1 cells (*blue*) with 15 minutes EdU treatment. **d**. Coefficient of Variation (y-axis) versus average reads per bin (x-axis) for all single pulse scEdU-seq cells. Each dot is a single cell and the top area between three dashed lines contain selected cells for subsequent analysis. **e**. Schematic representation computing S-phase progression. *1 -* example distribution of reads from 3 different single cells A, B and C. *2 -* Example of gaussian kernel smoothing of the reads from single cells A and B. *3 -* Computing pairwise overlap coefficient between all cells. *4 -* Converting overlap into distance metric between single cells**. f**. Dimensional of distance between single cells by UMAP. Each dot is a single cell, lines indicate nearest neighbors, dots are colored by S-phase progression, DNA content (DAPI) or FUCCI markers. **g**. Scatter plot showing FUCCI reporters versus DNA content. Dots are single cells pseudo colored by the S-phase progression based on scEdU-seq tracks and in gray the cell cycle distribution of cycling RPE-1 cells. **h**. Mid S-phase cells from scEdU-seq single pulse experiment for scRepli-seq comparison. DNA content (DAPI) intensities for Mid S-phase cells (blue) superimposed over cell cycle distribution of cycling RPE-1 cells (*grey*).

**Supplemental Figure 2:**
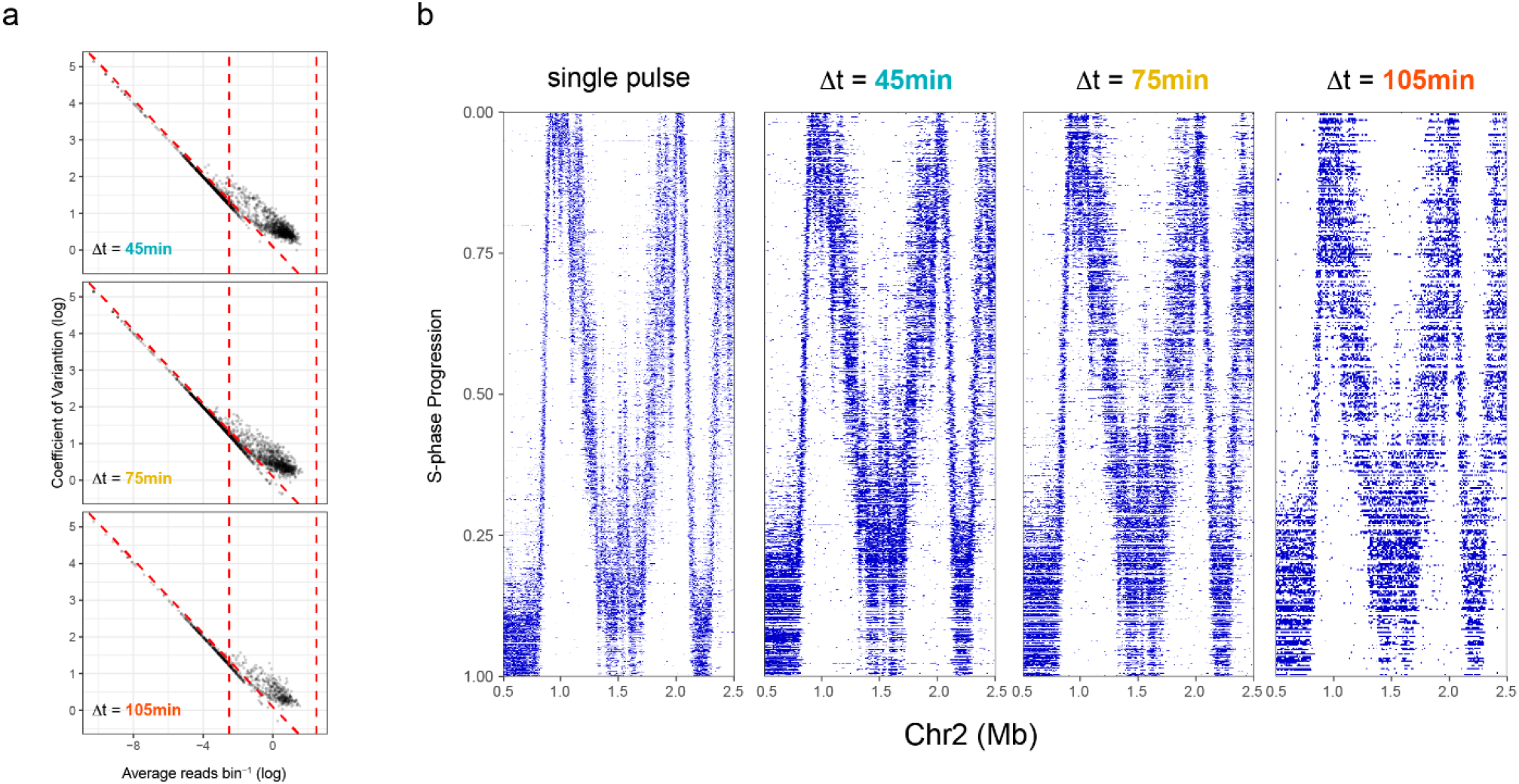
Supplemental Figure to Figure 2. **a**. Coefficient of Variation versus average reads per bin for all Δt=45min, Δt=75min and Δt=105min double pulse library scEdU-seq cells. Each dot is a single cell and the top area between three dashed lines contain selected cells for subsequent analysis. **b**. Heatmap of scEdU-seq maximum normalized log counts for all single pulse, Δt=45min, Δt=75min and Δt=105min double pulse cells ordered according to S-phase progression (*y-axis*) and binned per 5kb bins (x-axis) for 2 megabase of chromosome 2.

**Supplemental Figure 3:**
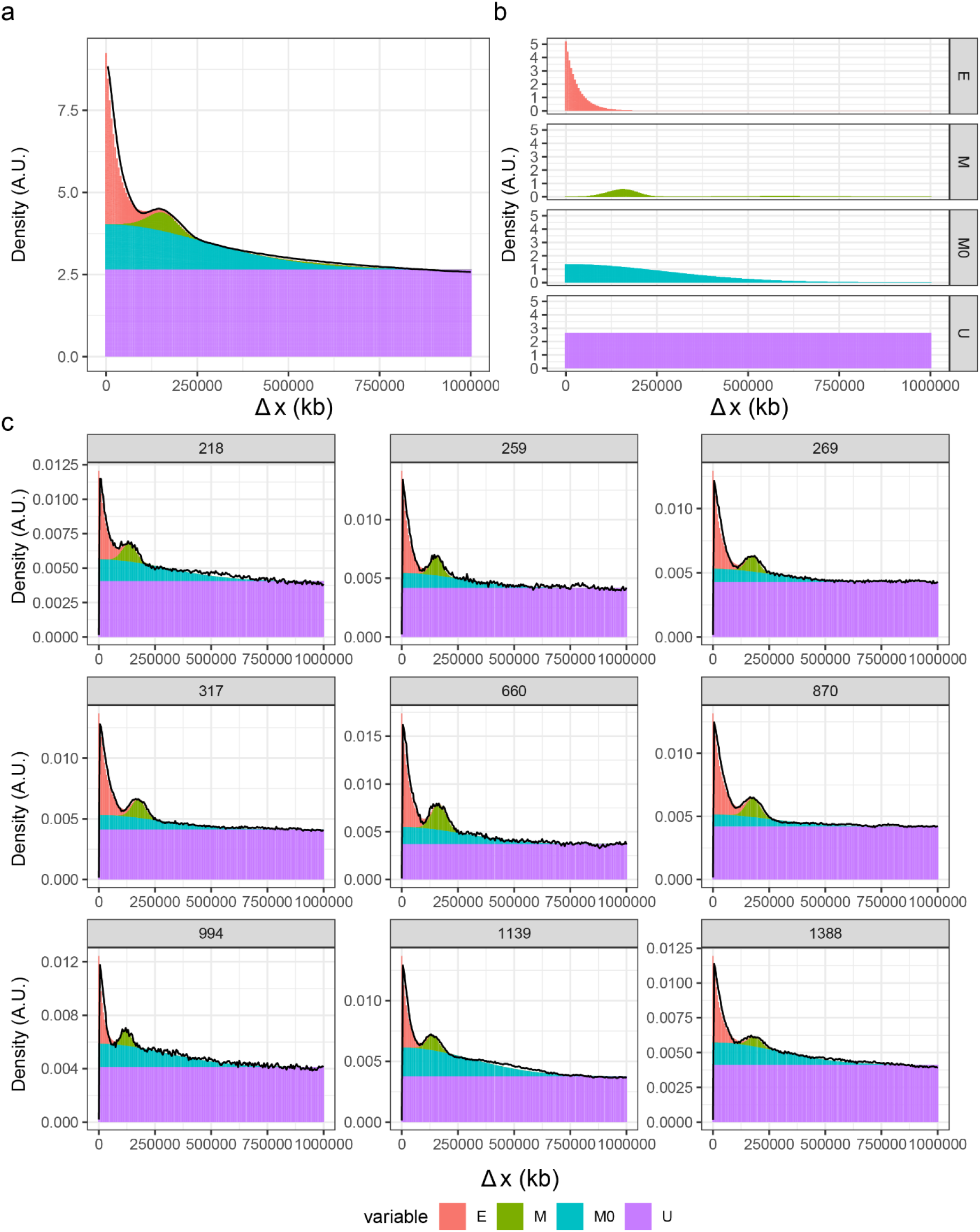
Supplemental Figure to Figure 3. **a**. Four component mixture model fit for all aggregate Δt=75min double pulse RPE-1 cells. Aggregate values per component per binned distance in kilobase (E= Exponent, N=normal (speed), M0 = half-normal, U= Uniform components). **b**. Four component mixture model fit for all aggregate Δt=75min double pulse RPE-1 cells split between all components. (E= Exponent, N=normal (speed), M0 = half-normal, U= Uniform components). **c**. Representative single cells with Four component mixture model fit for Δt=75min double pulse RPE-1 cells. (E= Exponent, N=normal (speed), M0 = half-normal, U= Uniform components).

**Supplemental Figure 4:**
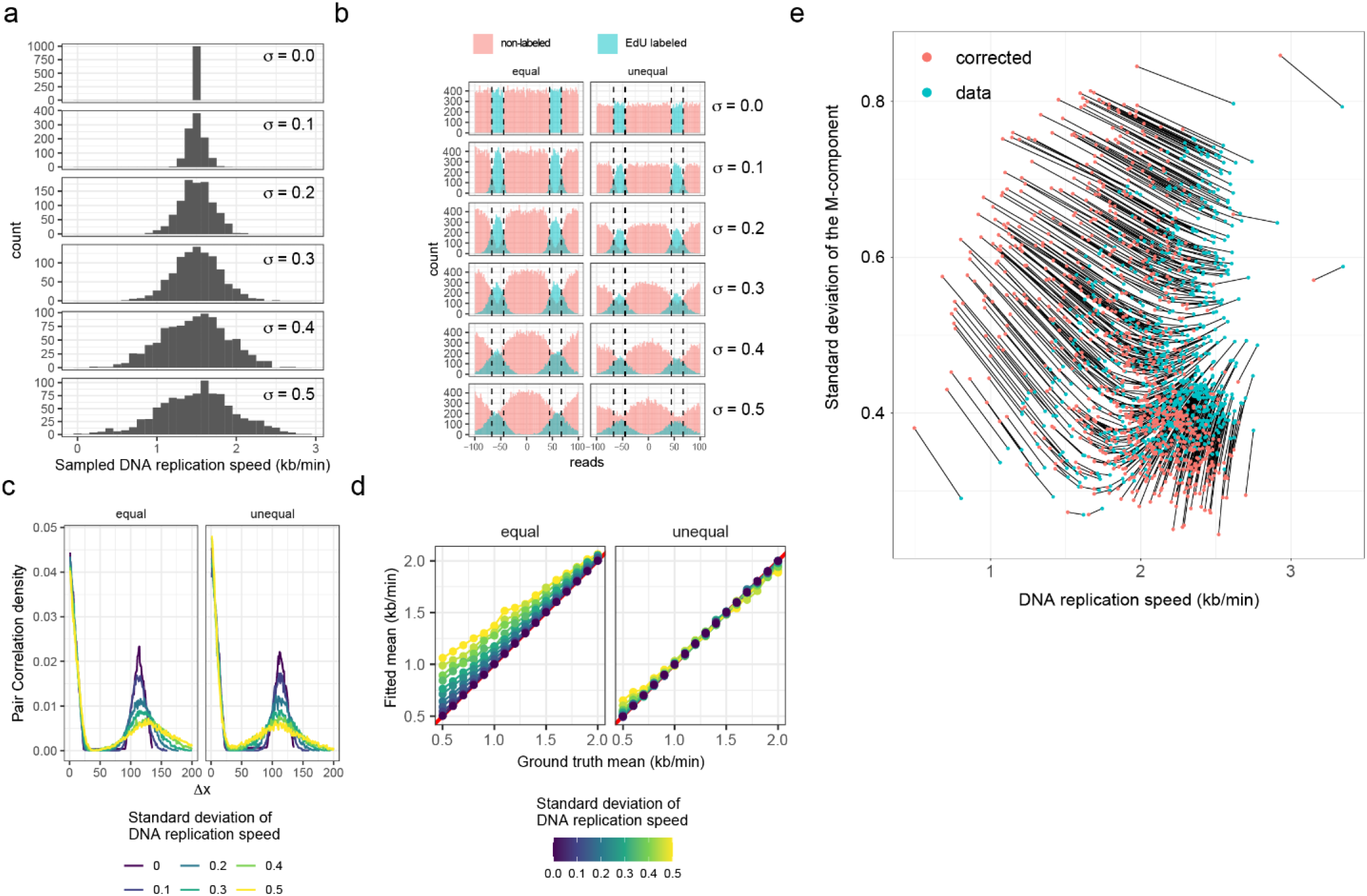
Supplemental FIgure to Figure 3. a. Simulated DNA replication speeds drawn from a truncated normal distribution with a mean of 1.5kb/min and increasing standard deviation (0-0.5kb/min). **b**. Density of reads (y-axis) aggregated per standard deviation of the underlying speed distribution for the piece of simulated genome (x-axis), colored by EdU (inside window) or non-labeled (outside window) and faceted by the poisson intensity assumption. **c**. Line plots of the pair correlation for simulated double pulse labeling for equal and unequal sampling with increasing DNA replication speed variance.The x-axis shows the binned distance and the y-axis the range-scaled density. Line colors indicate the underlying DNA replication speed variance. **c**. Average of the between-pulse-distances from the simulated data (y-axis) versus the ground-truth simulated mean (x-axis). Line colors indicate the ground-truth simulated mean variance and the facets show the equal and unequal sampling assumptions. **d**. Standard deviation (y-axis) versus the mean (x-axis) of the speed component where every dot is colored indicating whether the fitted or corrected values are plotted, and where dots representing the same cell before and after correction are connected by a line.

**Supplemental Figure 5:**
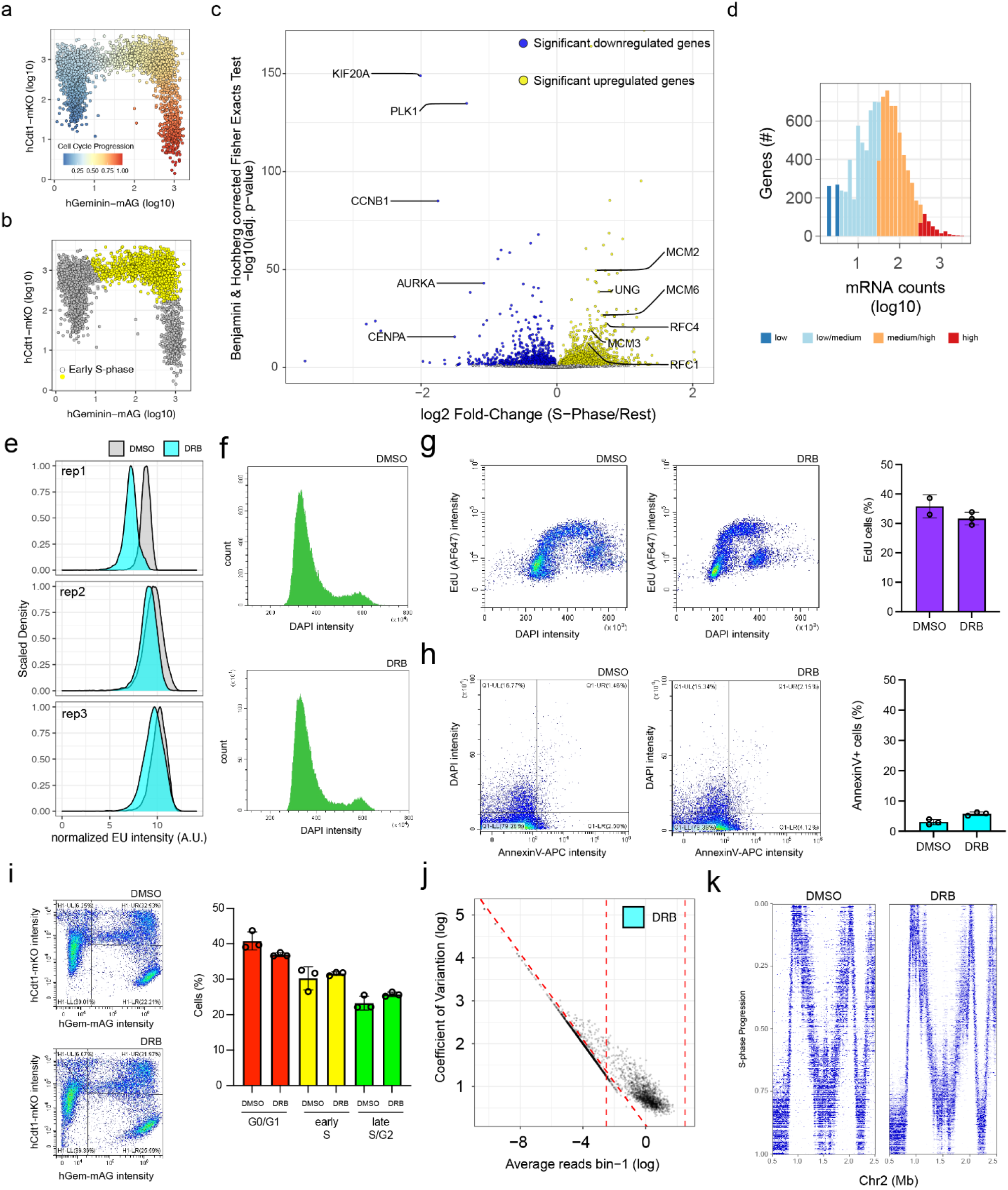
Supplemental Figure to Figure 3. **a**. FUCCI reporter intensities with pseudotime ordering (wanderlust) based on scEUseq counts (Battich et al., 2020(33)). **b**. Early S-phase cells selected for gene expression analysis from scEUseq data in Figure 3c. **c**. Differentially expressed genes between early S-phase cells versus the rest. Volcano plot displaying log2 Fold Change (x-axis) versus adjusted Fisher exact test (Benjamini-Hochberg correction for multiple testing). Each dot represents a gene and is colored by significance (adj. p-value <0.05) and up/down sign. **d**. Distribution of log10-transformed nascent mRNA counts for pseudo-bulked S-phase cells **e**. Nascent RNA labeling with EU (30min) on fixed RPE-1 cells treated with DMSO or DRB (1hr). **f**. Representative DNA content (DAPI) intensities for fixed RPE-1 cells treated with DMSO or DRB (1hr). **g**. Representative EdU (30min) intensities for fixed RPE-1 cells treated with DMSO or DRB for 1hr (*left*). Quantification of EdU^+^ cells, each dot indicates a biological replicate (*right*). **h**. Representative AnnexinV/DAPI intensities on living RPE-1 cells treated with DMSO or DRB for 1hr (*left*). Quantification of AnnexinV/DAPI^+^ cells, each dot indicates a biological replicate (*right*). **i**. Representative FUCCI reporter intensities on fixed RPE-1 cells treated with DMSO or DRB (*left*). Quantification of G0/G1, early S and late S-phase/G2 cells, each dot indicates a biological replicate (*right*). **j**. Coefficient of Variation (y-axis) versus average reads per bin (x-axis) for all DRB-treated Δt=75min scEdU-seq cells. Each dot is a single cell and the top area between three dashed lines contain selected cells for subsequent analysis. **k**. Heatmap of scEdU-seq maximum normalized log counts for DMSO and DRB-treated t=75min double pulse cells ordered according to S-phase progression (*y-axis*) and binned per 5kb bins (x-axis) for 2 megabase of chromosome 2.

**Supplemental Figure 6:**
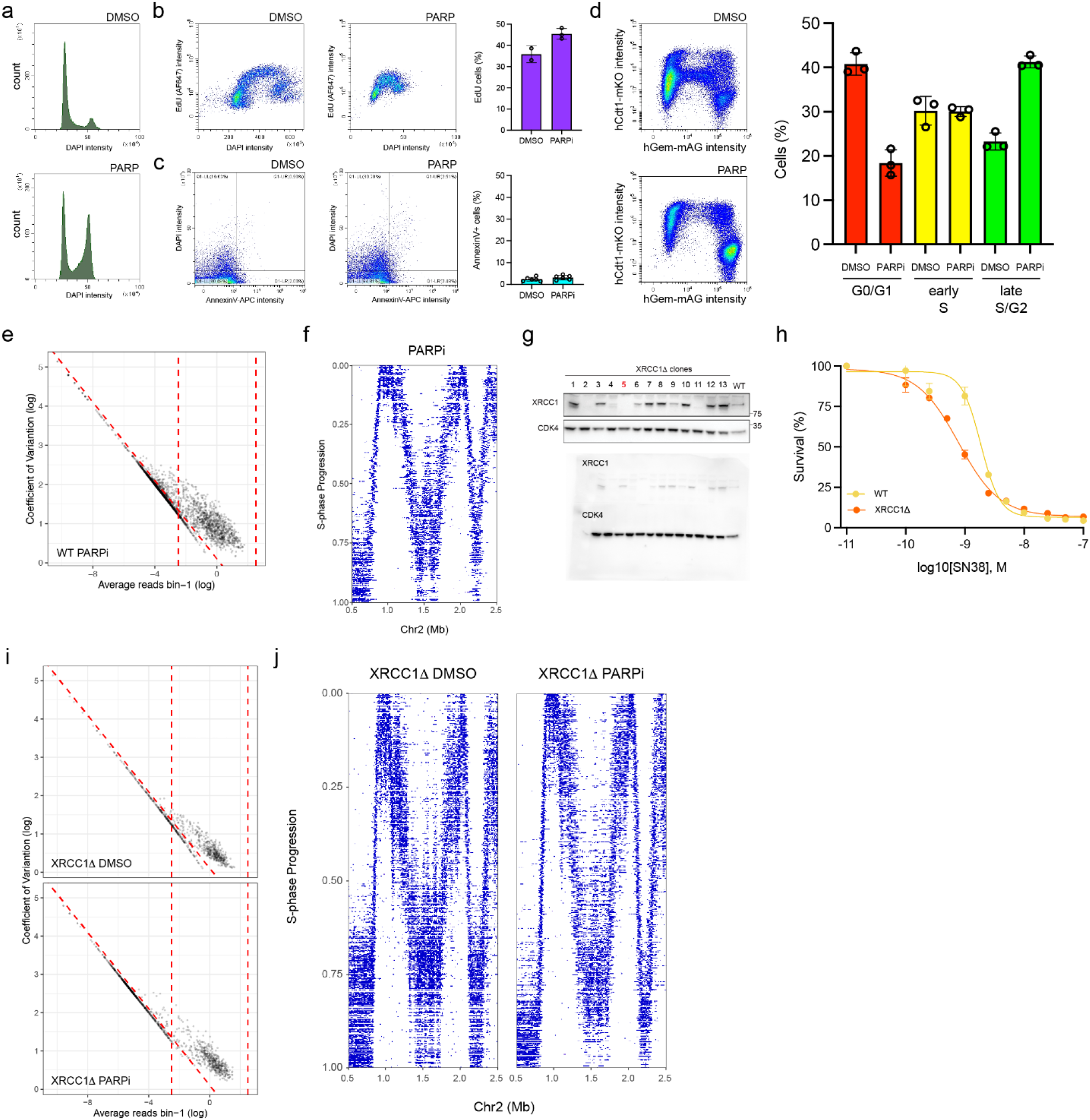
Supplemental Figure to Figure 4. **a**. Representative DNA content (DAPI) intensities for fixed RPE-1 cells treated with DMSO or 24 hr PARPi. **b**. Representative EdU (30min) intensities for fixed RPE-1 cells treated with DMSO or 24hr PARPi (*left*). Quantification of EdU^+^ cells, each dot indicates a biological replicate (*right*). **c**. Representative AnnexinV/DAPI intensities on living RPE-1 cells treated with DMSO or 24 hr PARPi (*left*). Quantification of AnnexinV/DAPI^+^ cells, each dot indicates a biological replicate (*right*). **d**. Representative FUCCI reporter intensities on fixed RPE-1 cells treated with DMSO or 24hr PARPi (*left*). Quantification of G0/G1, early S and late S-phase/G2 cells, each dot indicates a biological replicate (*right*). **e**. Coefficient of Variation (y-axis) versus average reads per bin (x-axis) for all 24hr PARP inhibitor treated Δt=75min scEdU-seq cells. Each dot is a single cell and the top area between three dashed lines contain selected cells for subsequent analysis. **f**. Heatmap of scEdU-seq maximum normalized log counts for PARP-treated t=75min double pulse cells ordered according to S-phase progression (*y-axis*) and binned per 5kb bins (x-axis) for 2 megabase of chromosome 2. **g**. Western Blot analysis of XRCC1Δ RPE-1 clones with XRCC1 antibody to validate knockout status and CDK4 as a loading control (*top*) and uncropped blots (*bottom*). **h**. Viability of XRCC1Δ RPE-1 and parental RPE-1 cells in response to increasing concentrations of Topoisomerase I poison, SN-38. **i**. Coefficient of Variation (y-axis) versus average reads per bin (x-axis) for all XRCC1Δ RPE-1 DMSO and 4hr PARPi-treated Δt=75min scEdU-seq cells. Each dot is a single cell and the top area between three dashed lines contain selected cells for subsequent analysis. **j**. Heatmap of scEdU-seq maximum normalized log counts for XRCC1Δ RPE-1 DMSO and PARPi-treated Δt=75min scEdU-seq cells ordered according to S-phase progression (*y-axis*) and binned per 5kb bins (x-axis) for 2 megabases of chromosome 2.

